# Enhancing Gesture Decoding Performance Using Signals from Posterior Parietal Cortex: A Stereo-Electroencephalograhy (SEEG) Study

**DOI:** 10.1101/849752

**Authors:** Meng Wang, Guangye Li, Shize Jiang, Zixuan Wei, Jie Hu, Liang Chen, Dingguo Zhang

## Abstract

**Objective:** Hand movement is a crucial function for humans’ daily life. Developing brain-machine interface (BMI) to control a robotic hand by brain signals would help the severely paralyzed people partially regain the functional independence. Previous intracranial electroencephalography (iEEG)-based BMIs towards gesture decoding mostly used neural signals from the primary sensorimotor cortex while ignoring the hand movement related signals from posterior parietal cortex (PPC). Here, we propose combining iEEG recordings from PPC with that from primary sensorimotor cortex to enhance the gesture decoding performance of iEEG-based BMI.

**Approach:** Stereoelectroencephalography (SEEG) signals from 25 epilepsy subjects were recorded when they performed a three-class hand gesture task. Across all 25 subjects, we identified 524, 114 and 221 electrodes from three regions of interest (ROIs), including PPC, postcentral cortex (POC) and precentral cortex (PRC), respectively. Based on the time-varying high gamma power (55-150 Hz) of SEEG signal, both the general activation in the task and the fine selectivity to gestures of each electrode in these ROIs along time was evaluated by the coefficient of determination *r*^2^. According to the activation along time, we further assessed the first activation time of each ROI. Finally, the decoding accuracy for gestures was obtained by linear support vector machine classifier to comparatively explore if the PPC will assist PRC and POC for gesture decoding.

**Main Results:** We find that a majority(*L: >*60%, *R: >*40%) of electrodes in all the three ROIs present significant activation during the task. A large scale temporal activation sequence exists among the ROIs, where PPC activates first, PRC second and POC last. Among the activated electrodes, 15% (PRC), 26% (POC) and 4% (left PPC) of electrodes are significantly selective to gestures. Moreover, decoding accuracy obtained by combining the selective electrodes from three ROIs together is 5%, 3.6%, and 8% higher than that from only PRC and POC when decoding features across, before, and after the movement onset, were used.

**Significance:** This is the first human iEEG study demonstrating that PPC contains neural information about fine hand movement, supporting the role of PPC in hand shape encoding. Combining PPC with primary sensorimotor cortex can provide more information to improve the gesture decoding performance. Our results suggest that PPC could be a rich neural source for iEEG-based BMI. Our findings also demonstrate the early involvement of human PPC in visuomotor task and thus may provide additional implications for further scientific research and BMI applications.

## 1. Introduction

For the severely paralyzed people, brain-machine interface (BMI) is a promising solution towards helping them regaining the sensorimotor interaction with the external physical world via reading brain signals and converting it into commands to control external devices (Collinger et al., 2013). For BMI purpose, human intracranial electroencephalography (iEEG) recordings (including electrocorticography (ECoG) and stereoelectroencephalography (SEEG), (Parvizi and Kastner, 2018)), holds the advantage of stably recording rich neural information across multiple brain regions. Thus, iEEG-based BMIs have made significant achievement in reconstructing three-dimensional reach function (Talakoub et al., 2017; Hu et al., 2018; Nakanishi et al., 2017; Flint et al., 2014; Wang et al., 2013; Yanagisawa et al., 2012; Fifer et al., 2014). But for the equally crucial hand movement decoding, although several iEEG-based studies have succeeded in one-dimensional hand kinematics regression or individual finger classification (Kubanek et al., 2009; Acharya et al., 2010; Flint et al., 2014; Nakanishi et al., 2014; Hotson et al., 2016), for functional use there is still a wide gap to bridge because it requires multi-finger coordination at much higher dimensions. To facilitate this, other studies decode frequently-used functional gestures directly as an alternative solution and have achieved considerable decoding accuracy (Pistohl et al., 2012; Chestek et al., 2013; Bleichner et al., 2014; Spueler et al., 2014; Branco et al., 2017; Li et al., 2017). Nevertheless, most of these iEEG studies mainly focus neural signals from primary sensorimotor cortex (i.e., precentral and postcentral cortex, abbreviated as PRC and POC respectively), to decode gestures, which works well but may not optimally.

Beyond primary sensorimotor cortex, posterior parietal cortex (PPC) has been reported to be related with multiple hand movements like grasp, intransitive postures and even pantomime and imagination of those movements (Culham and Valyear, 2006; Vingerhoets, 2014), and moreover, contribute to the formation of early motor intention (Andersen and Buneo, 2002). Lesion of PPC causes severe deficits of controlling hand shape appropriate for object (Jeannerod et al., 1994). Intracortical electrical stimulation to lateral region of monkey’s area 5 within PPC elicits finger and wrist movements (Rathelot et al., 2017). Functional magnetic resonance imaging (fMRI) studies reveal that grip type (power vs precision grasp) is encoded in several subareas of PPC (Fabbri et al., 2016; Di Bono et al., 2015). In addition to object-related grasp, other intransitive hand gestures like scissor-rock-paper can also be decoded from PPC, as demonstrated by another fMRI study (Dinstein et al., 2008). Electrophysiological studies on macaque also demonstrate that some neurons in the anterior intraparietal sulcus (AIP), as a subarea of PPC, are tuned to grip type (Baumann et al., 2009). It allows the real-time prediction of grip type (Townsend et al., 2011) by spiking activity with an accuracy of 90%. Beyond grip type, Menz et al. (2015) reconstructs 27 degrees of freedom representing continuous hand and arm kinematics using spiking activity in AIP of PPC with a correlation coefficient about 0.5 between the reconstructed and actual kinematics, demonstrating a substantial amount of temporal kinematic information in PPC. In further human study on tetraplegic patient by Klaes et al. (2015), the imagination of scissor-rock-paper hand gestures can be decoded by spiking activity in PPC. Taken together, all these evidences support the notion that PPC has distinct relevance to hand movement control.

However, although electrophysiological studies have demonstrated that spiking activity from PPC is involved in hand shape encoding, whether iEEG recordings from human PPC could also benefit gesture decoding is still unclear. There are some promising clues. (1) Previous macaque local field potential (LFP) studies (Pesaran et al., 2002; Scherberger et al., 2005) have shown that high-frequency LFP in PPC is coherent to spiking. The LFP signals are also tuned to direction of reach and saccade, and thus manifest as good decoding performance as spiking activity; (2) Asher et al. (2007) recorded LFP and spiking activity simultaneously from several areas of macaque PPC during grasp prehension and find that LFP presents selectivity to different grasp prehensions as well in a less informative way than spiking; (3) In two human ECoG study towards hand movement decoding, some electrodes in PPC present high decoding accuracies (Pistohl et al., 2012; Chestek et al., 2013) but they were not analyzed in detail. Accordingly, we speculate that human iEEG recordings from PPC contains rich information about hand gestures and could further play a complementary role to primary sensorimotor cortex in enhancing of the performance of iEEG based BMI.

To justify the hypothesis, we record signals from 25 human SEEG subjects performing a hand gesture task, where the subjects are implanted with multiple depth electrodes (Bartolomei et al., 2018; Parvizi and Kastner, 2018; Arnal et al., 2019). SEEG has the ability of recording across multiple brain sites simultaneously with high temporal resolution. Therefore, SEEG recordings across subjects allows an exhaustive investigation on a wide human PPC area about its involvement in hand motor task. Beyond wide coverage, SEEG electrodes can reach both the gyri and sulci of PPC (e.g., the intraparietal sulcus (IPS), which is important for hand movement (Sakata et al., 1995; Fabbri et al., 2016)). Using the SEEG recordings, we first investigate whether and how much the neural activities, recorded from electrodes that distributed in PPC and primary sensorimotor cortex (PRC and POC), contain activation information relating to the task and selective information with respect to specific gestures. We then assess at what time the neural signals from these areas get activated during the task. We observe a relatively small group of PPC channels contain gesture selective information while a majority of them present task related activation and they tend to activate earlier than the primary sensorimotor cortex. On the basis of our findings, we combine the neural signals from selective electrodes in PPC and primary sensorimotor cortex to decode the gestures. The results show that the introduction of PPC improves decoding accuracy.

## 2. Methods

### 2.1. Subject

Twenty five subjects in total participated in the study. These subjects were intractable epilepsy patients under pre-surgical assessment of their seizure focus and SEEG electrodes were implanted for this purpose (See table 1 for more details). Each implanted electrode shaft was 0.8 mm in diameter and contained 8-16 contacts along the length direction. Each contact was 2 mm long and spaced at 3.5 mm center-to-center. The entire implantation was solely determined by clinical needs. The study was approved by Ethics Committee of Huashan Hospital (Shanghai, China). All subjects were fully informed and signed written consent.

**Table 1:** Subject information of this study. Abbreviation List: Subj, Subject; TH, Task Hand; EH, Electrode Implantation Hemisphere; NS, Number of Electrode Shafts; NE, Number of Electrode Contacts; SR, Sampling Rate. Notes: N/A in Seizure Focus column indicates the seizure focus is unknown because either the seizure symptom is not ameliorated after surgery or the resection surgery is not performed. B in EH column indicates the electrodes are implanted bilaterally. All subjects report right handedness.

| Subj | Age | Gender | TH | Seizure Focus | EH | NS | NE | SR (Hz) |
| --- | --- | --- | --- | --- | --- | --- | --- | --- |
| S1 | 33 | M | R | left occipital | L | 15 | 180 | 1000 |
| S2 | 30 | F | L | right frontal | R | 7 | 60 | 1000 |
| S3 | 27 | M | R | left temporal | L | 16 | 208 | 2000 |
| S4 | 22 | F | R | left frontal | L | 7 | 102 | 2000 |
| S5 | 24 | M | L | right temporal | R | 8 | 108 | 1000 |
| S6 | 23 | M | R | left frontal | L | 10 | 121 | 1000 |
| S7 | 26 | M | L | right temporal | R | 13 | 178 | 1000 |
| S8 | 25 | M | L | right frontal | R | 10 | 143 | 1000 |
| S9 | 17 | F | R | right temporal | B | 13 | 169 | 1000 |
| S10 | 28 | F | L | right frontal | R | 9 | 114 | 1000 |
| S11 | 15 | M | L | right temporal | B | 13 | 194 | 500 |
| S12 | 31 | M | L | right parietal | R | 6 | 94 | 500 |
| S13 | 19 | M | L | right cingulate | B | 9 | 130 | 2000 |
| S14 | 30 | F | R | left insula | B | 13 | 170 | 2000 |
| S15 | 31 | M | R | left temporal | L | 10 | 144 | 2000 |
| S16 | 27 | M | R | left occipital | B | 10 | 144 | 2000 |
| S17 | 16 | M | R | N/A | B | 13 | 137 | 2000 |
| S18 | 30 | F | R | left temporal | L | 9 | 118 | 1000 |
| S19 | 33 | F | R | left temporal | L | 12 | 150 | 2000 |
| S20 | 23 | F | R | N/A | B | 15 | 198 | 2000 |
| S21 | 23 | F | L | right temporal | R | 10 | 130 | 2000 |
| S22 | 42 | F | R | left temporal | L | 10 | 137 | 2000 |
| S23 | 33 | M | R | left temporal | B | 11 | 154 | 2000 |
| S24 | 15 | M | R | left frontal | L | 8 | 110 | 2000 |
| S25 | 25 | M | L | right occipital | B | 8 | 107 | 2000 |

### 2.2. Data Recording

Neural signals were recorded using a clinical recording system (EEG-1200C, Nihon Kohden, Irvine, CA) with a sampling rate ranged from 500 to 2000 Hz (See table 1). Electromyography (EMG) signals were acquired by two surface electrodes placed on extensor carpiradialis muscle from the same recording system with SEEG signals simultaneously.

### 2.3. Task Protocol

The task protocol is illustrated in figure 1. In detail, the subjects were instructed by the picture displayed on a LCD screen to perform three different gestures, i.e., scissor, rock and thumb. Each trial of the main task lasted 10 s and consisted of warning (1 s) task (5 s) and rest (4 s) period. During the warning period, a cross was first shown to remind the upcoming hand movement task. Then, during the task period, one of three gestures, including scissor, rock and thumb, was displayed as the Cue (figure 1 (B)). The subjects repeatedly performed the hand movement following the Cue using the hand contralateral to the hemisphere with the majority of the implanted SEEG electrodes (see task hand in table 1). During the rest period, the subjects kept relaxed without any hand movement. 20 trials were performed for each gesture. Overall, the task consisted of 60 trials and lasted about 10 minutes in total per subject.

**Figure 1:**
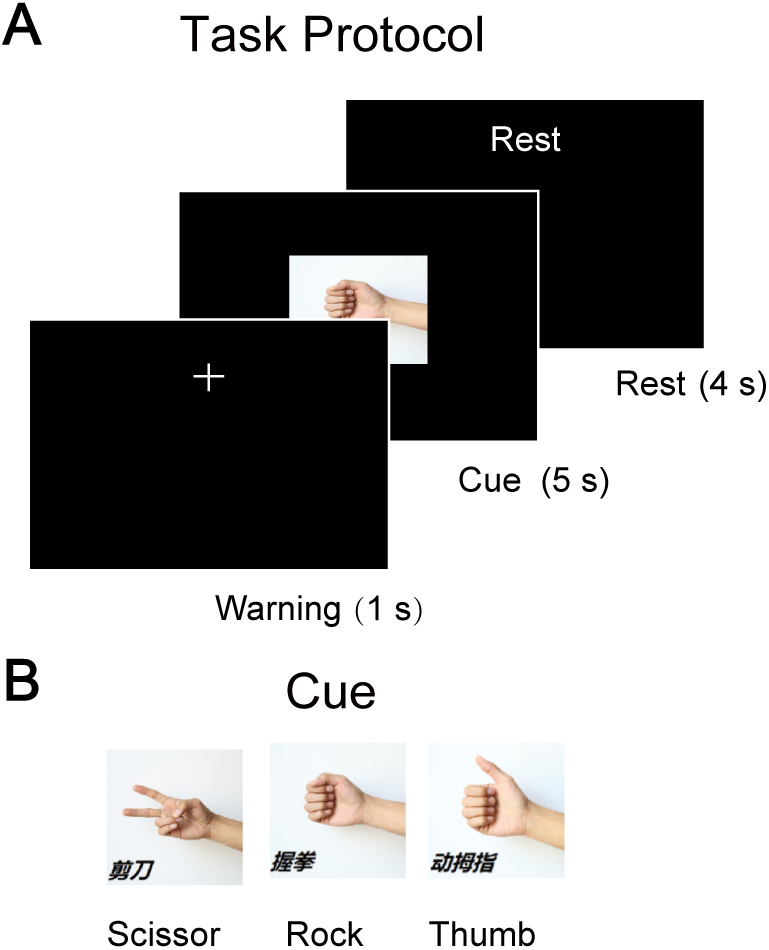
Task protocol. **(A)** The protocol started from a 1-s warning phase to alert the subject. Next, a gesture picture appeared for 5 s (Cue phase) to instruct the subject to repeatedly perform the corresponding gesture. Following that was a 4-s rest phase, the subject kept relaxed without any movement. 20 trials for each gesture were performed for each subject. **(B)** The possible gesture pictures (i.e., scissor, rock and thumb) shown in the Cue phase of (A). For each trial, one of them was randomly selected while ensuring an equal number of appearances.

### 2.4. Electrode localization

Each electrode was localized in the individual brain model according to the following steps. First, the individual brain model was reconstructed and segmented using pre-implantation MRI images through Freesurfer (http://surfer.nmr.mgh.harvard.edu). Second, the coordinates of each electrode contact was determined by co-registering post-implantation CT images with pre-implantation MRI images. The anatomical label of each electrode contact was identified by cortical parcellation and subcortical segmentation results under the Desikan-Killiany atlas (Desikan et al., 2006; Fischl et al., 2002). Finally, each electrode was mapped from the individual brain to a standard MNI (Montreal Neurological Institute) brain template for the purpose of group analysis. Since we were interested in the neural response in PRC, POC and PPC, therefore, only electrodes within these regions of interest (ROIs) were included in this study. Specifically, PPC was divided by IPS into superior parietal lobe (SPL) and inferior parietal lobule (IPL). IPL consisted of supramarginal gyrus (SMG) and angular gyrus. In the parietl lobe of Desikan-Killiany atlas, they were corresponding to superior parietal cortex, supramarginal gyrus, and inferior parietal cortex, respectively. Electrodes located at white matter were identified using white matter segmentation and were included in the study as well, considering the observations that white matter presented similar neural activities with gray matter under task (Ding et al., 2018; Mercier et al., 2017). All the procedures above were implemented with iEEGview toolbox (Li et al., 2019). Additionally, the anatomical position of all electrodes located in ROIs were visually inspected by experienced neurosurgeons. The final distribution of the valid electrodes used in this work was shown in figure 2.

**Figure 2:**
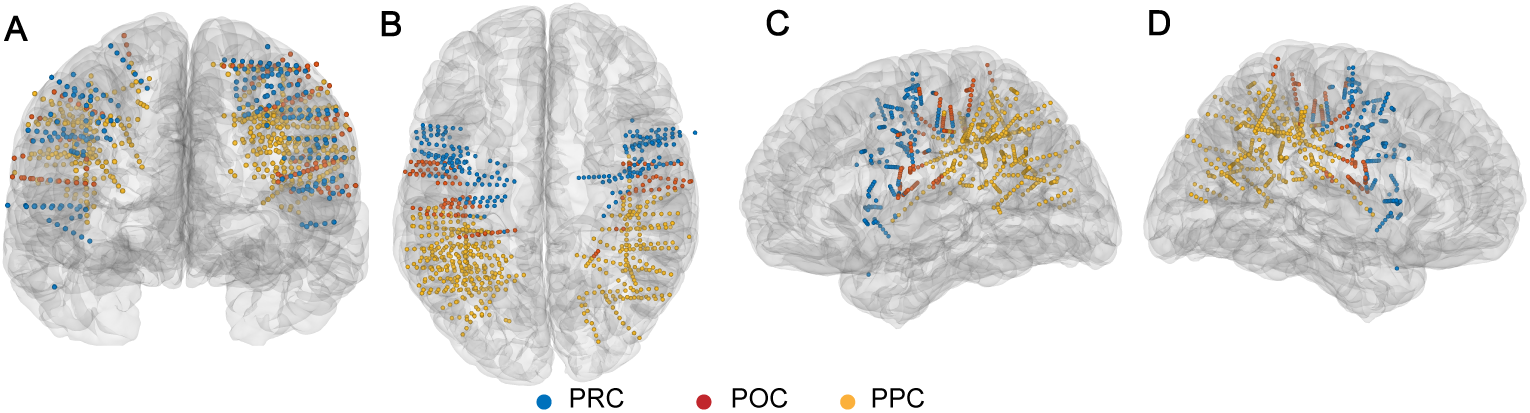
**A/B/C/D)** Front/Top/Left/Right view of the locations of SEEG electrodes in a standard MNI (Montreal Neurological Institute) brain template. All electrodes within ROIs from 25 subjects are shown. Each colored dot represents one electrode contact in a specific ROI. *Blue*: precentral cortex (PRC), *red*: postcentral cortex (POC), *golden*: posterior parietal cortex (PPC).

To visualize the activation, we implemented a cross-subject mapping on the inflated standard brain surface using the “Activation Map Visualization” function from the iEEGview toolbox. Specifically, the mapping strength on each vertex of brain was the weighted sum of *r*^2^ value (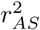 in Sec. 2.6.1 and 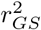 in Sec. 2.6.2) of each electrode.

### 2.5. Data Processing

All the recorded SEEG data were processed using custom Matlab (MathWorks, Natick, MA, USA) scripts according to following pipeline. The raw signals were first resampled to 1000 Hz uniformly to reduce computational cost and facilitate further computation. Then, the channels whose line noise power at 50 Hz were larger than a significance level were removed from further analysis. The significance level was defined as the summation of median line noise power across all electrodes and 10 times of their median absolute deviation. At the third step, all the process signals were subjected to a 50 Hz comb notch filter to remove the possible line noises and its harmonics. After this, the signals were re-referenced using a Laplacian scheme to further improve the signal quality (Li et al., 2018). In brief, the signals of each time point were subtracted by the average signals from adjacent channels at the same time point. We then extracted the high gamma power (HGP) from the processed signals, since the high gamma activity had been proven to contain the richest gesture information (Asher et al., 2007; Bleichner et al., 2014). In detail, the re-referenced signals were first band-pass filtered at [55, 150] Hz using sixth order Butterworth filter and then the HGP were extracted by computing the squared absolute value of Hilbert-transformed signals at this frequency band. At the next step, we aligned HGP signals to the movement onset for each trial, where the movement onset was defined as the time point when the absolute amplitude of EMG first time exceeded an adaptive threshold using the methods described in (Sedghamiz, 2018). To be concise, the movement onset was denoted as *Move* and the time when the Cue (figure 1) appeared was denoted as *Cue* throughout the following article. The median reaction time was 564 ms across 25 subjects (see supplementary Figure 1 for the detected *Move* time information of each subject). Afterwards, a two-step normalization was applied to the HGP of each electrode. First, the HGP during task subtracted baseline within each trial to extract the change during task. Second, the HGP across all the trials of each electrode were concatenated and zscored to eliminate the difference of absolute amplitude among electrodes. The baseline period was defined as the [−0.75, −0.25] s around the *Cue*. At the final step, the normalized HGP signals were averaged every non-overlapping 100 ms window, and thus generating a HGP feature vector with a dimension of *NW × NE* per trial for each subject (*NW*, number of windows; *NE*, number of electrodes). The derived HGP feature vector was further used in the following performance index calculation. Due to windowing, the continuous time course became discrete, therefore, we referred the time of each window as its middle time point for convenience. For instance, HGP at 0.15 s represented the average HGP at [0.1, 0.2] s.

### 2.6. Performance Index

To evaluate the characteristics of neural response and possible functions from these ROIs (PRC, POC and PPC) in relation to the hand movement, we proposed four performance indices in this study: 1) Activation strength (AS); 2) Gesture selectivity (GS); 3) First activation time (FAT); and 4) Decoding accuracy (DA).

#### 2.6.1. Activation Strength

To evaluate the engagement of neural activities relating to the task, we computed the activation strength (AS) using the coefficient of determination *r*^2^ (equation 1) (Pfurtscheller et al., 2006; Kubanek et al., 2009), denoted as 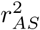 in this work. It measures how much total variance of HGP could be explained by the difference between task and rest. 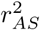 was calculated at each electrode and each time window to evaluate the neural response at a temporal and spatial scale. In detail, in equation 1, *X*_*i,j*_ denoted the HGP of class *i* (*i* = 1…*c, c* = 2, i.e., rest or task state) at *j*^*th*^ trial (*j* = 1*…m, m* = 60), 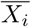 denoted the mean HGP across trials of class *i* and 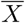 denoted the mean HGP across classes and trials. We then implemented a permutation test (Edgington, 1998) with 3000 repeats to determine the significantly activated electrodes across all electrodes in each subject, where the label (i.e., rest and task) was randomly permuted across trials and corresponding *r*^2^ was computed based on the permuted label, thereby generating null distribution of *r*^2^ and revealing *p* value of observed *r*^2^. The *p* value yielded by permutation test was FDR corrected for each subject by a factor of *NW × NE* (length of the feature vector, see Sec.2.5) (Storey and Tibshirani, 2003) due to multiple comparison. Any electrode, whose *p* value obtained after correction was smaller than 0.001 at any time window within [-0.5, 0.5] s around *Move*, was referred as activated electrode throughout the paper.

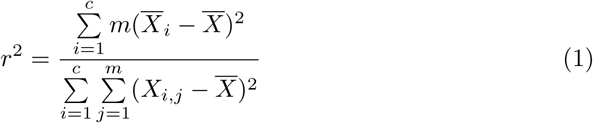

#### 2.6.2. Gesture Selectivity

To measure the difference of neural response to gestures within each ROI, we then calculated the gesture selectivity (GS) using a similar procedure with AS (Sec. 2.6.1), denoted as 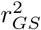. The only difference between AS and GS was that 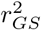 treated the gestures (*k* = 3, i.e., scissor, rock and thumb) as classes instead of the movement state (*k* = 2, i.e., task and rest) used in 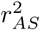 (equation 1, Sec. 2.6.1). It measures how much variance of HGP across all trials could be accounted for by the inherent deviation of classes from overall average. After this, random permutation test (3000 times) was conducted followed by FDR correction to identify the significantly selective electrodes of each subject (Sec.. 2.6.1). Any electrode, whose *p* value obtained here was smaller than 0.05 at any time window within [-0.5, 0.5] s around *Move*, was referred as selective electrode throughout the paper.

#### 2.6.3. First Activation Time

The first activation time was computed to determine when the neural activity around an SEEG electrode started to involving in the task significantly. In detail, FAT of an electrode was defined as the first time when 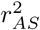 during [-0.5, 0.5] s around *Move* presented significance (*p <* 0.001, Sec. 2.6.1). Next, by a bootstrapping analysis with 1000 times randomized sampling, the 95% confidence interval of FAT was acquired. Furthermore, FAT was used to investigate the temporal activation sequence among the ROIs.

#### 2.6.4. Decoding Accuracy

To further assess the decoding performance for the ROIs and to answer the question that whether the PPC could assist PRC and POC for gesture decoding, we finally conducted a gesture decoding process using the HGP features. In detail, we first included the HGP at the time windows spaced every 200 ms within [-0.5, 0.5] s around *Move* (Sec. 2.6.2) of each selective electrode into feature vector in a concatenated way, resulting in a feature vector with dimensions at five times the number of selective electrodes for each subject. Next, due to the limited samples in current study, forward optimal feature selection (fOFS) was applied to reduce the feature dimension and avoid overfitting. Specifically, we successionally selected the best one feature from the original feature vector and included it into the optimal feature subset until the cross-validated decoding accuracy (DA) stopped increased. The selection criterion of the best one feature was that the feature could, together with existing optimal feature subset, maximize the DA. Then, support vector machine (SVM) with a linear kernel setting was adopted for classification (Chang and Lin, 2011) and the mean accuracy after five-fold cross validation was obtained as the optimal DA for one fOFS procedure. During the cross validation of fOFS, the training and testing samples were randomly selected, which caused uncertainty of the optimal feature set. Hence, the fOFS procedure was repeated 100 times, generating 100 optimal feature subsets, and consequently, producing 100 optimal DAs in total. We then averaged these DAs to yield the final DA for each subject, which was used through the subsequent analysis. The standard error of these DAs was also obtained for each subject.

## 3. Results

### 3.1. The statistics of AS and GS within ROIs

The statistics of activated electrodes and selective electrodes within each ROI was presented in table 2. Across 25 subjects, 859 (L=522, R=337) SEEG electrodes within ROIs were collected in total. Specifically, there were 26% (n=221, L=122, R=99) contacts in PRC, 13% (n=114, L=49, R=39) contacts in POC, and 61% (n=524, L=325, R=199) contacts in PPC. The electrodes within each ROI at either hemisphere came from 44% (n=11) subjects on average. Among these contacts, 67% (n=147, L=93, R=54), 60% (n=68, L=49, R=19) and 51% (n=265, L=200, R=65) got activated during task in PRC, POC and PPC respectively, indicating a majority of electrodes within all three ROIs were involved in the task. Moreover, among the activated contacts, 15% (PRC, bilateral average), 26% (POC, bilateral average) and 4% (PPC, left) presented significant selectivity for gestures, indicating that a relative small group of electrodes were selective among the activated electrodes for PRC and POC, while a minority for PPC.

**Table 2:**
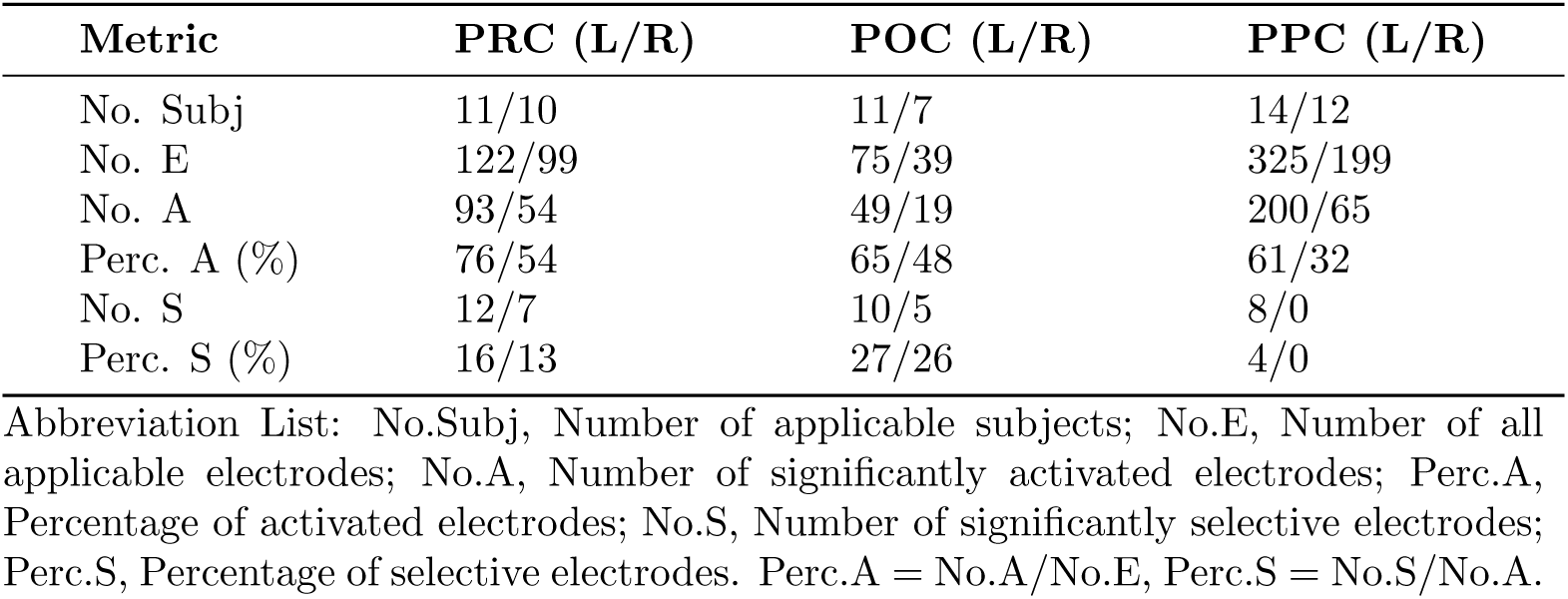
Response statistics of electrodes within ROIs from left and right (L/R) hemisphere.

| Metric | PRC (L/R) | POC (L/R) | PPC (L/R) |
| --- | --- | --- | --- |
| No. Subj | 11/10 | 11/7 | 14/12 |
| No. E | 122/99 | 75/39 | 325/199 |
| No. A | 93/54 | 49/19 | 200/65 |
| Perc. A (%) | 76/54 | 65/48 | 61/32 |
| No. S | 12/7 | 10/5 | 8/0 |
| Perc. S (%) | 16/13 | 27/26 | 4/0 |

Abbreviation List: No.Subj, Number of applicable subjects; No.E, Number of all applicable electrodes; No.A, Number of significantly activated electrodes; Perc.A, Percentage of activated electrodes; No.S, Number of significantly selective electrodes; Perc.S, Percentage of selective electrodes. Perc.A = No.A/No.E, Perc.S = No.S/No.A.

To give a more direct sense on neural activity distribution in relation to activation and selectivity during the task from ROIs, we further computed the activation and selectivity map on a standard brain template (Sec. 2.4). The spatial distribution of activated and selective areas were illustrated in figure 3. As could be seen from figure 3, all ROIs presented activation and selectivity in response to the task. The neural activation was distributed in a wide area with strong amplitude within each ROI (figure 3A), while selectivity was relatively weakly presented on a small area (figure 3B) especially for PPC, indicating the subtle representation of finer hand movement.

**Figure 3:**
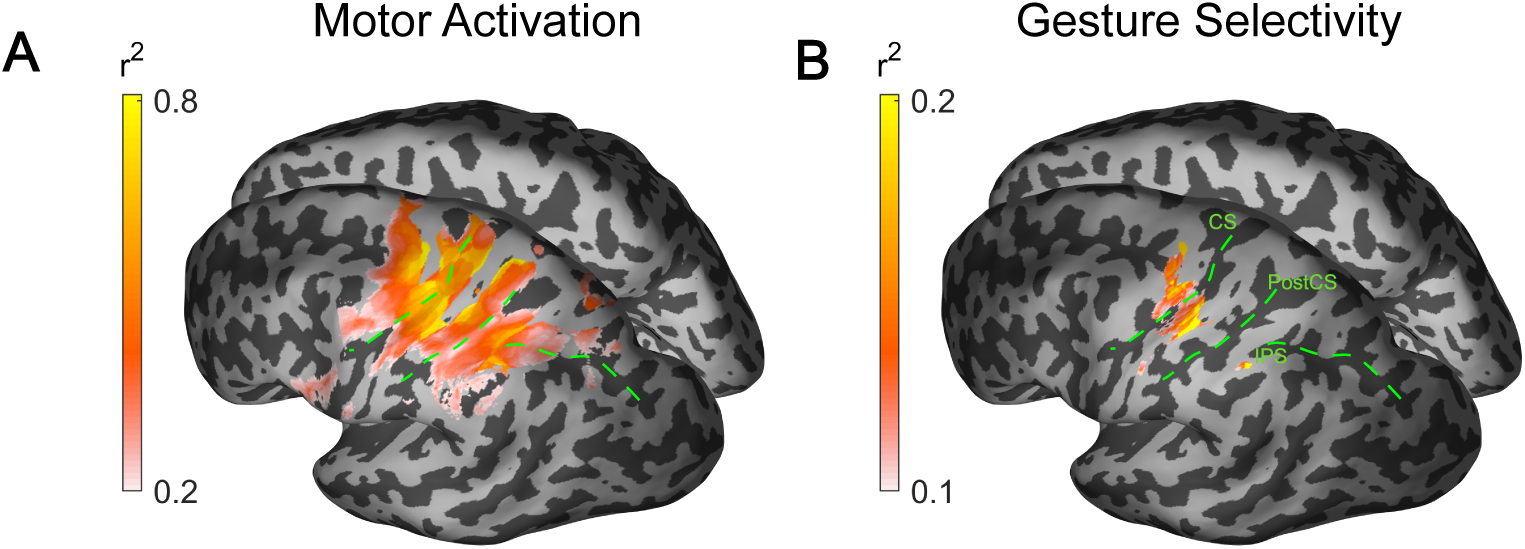
Spatial mapping of activation strength and gesture selectivity. For illustration purpose, the electrodes at right hemisphere are projected to its left counterpart. **(A)** Activation strength map. For each activated electrodes (see table 2), maximum 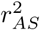 over time (see Sec. 2.6.1) is obtained as the activation value to compute the activation map. The projection is implemented on a standard inflated MNI brain template (for methods see Sec. 2.4). **(B)** Gesture selectivity map. The process is similar to (A), but maximum 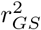 over time (see Sec. 2.6.2) is obtained as the selectivity value for each significantly selective electrodes to compute the selectivity map. The green dash line marks the three landmark sulci. CS, central sulcus; PostCS, postcentral sulcus; IPS, intraparietal sulcus.

### 3.2. The temporal activation sequence of ROIs

The response of electrodes from different ROIs had different characteristic patterns. As presented in figure 4A-B, the temporal HGP from an electrode located at PPC ascended immediately after *Cue* and reached summit before *Move*, finally returned to baseline after hundreds of milliseconds. The whole response lasted about 1.5 s. In contrast, as shown in figure 4C and D, HGP from the electrode near central sulcus climbed up to peak around *Move*, plateaued during movement execution, and finally declined to baseline after rest cue. The activation lasted about 6 s (See supplementary Figure 2 for another sample).

**Figure 4:**
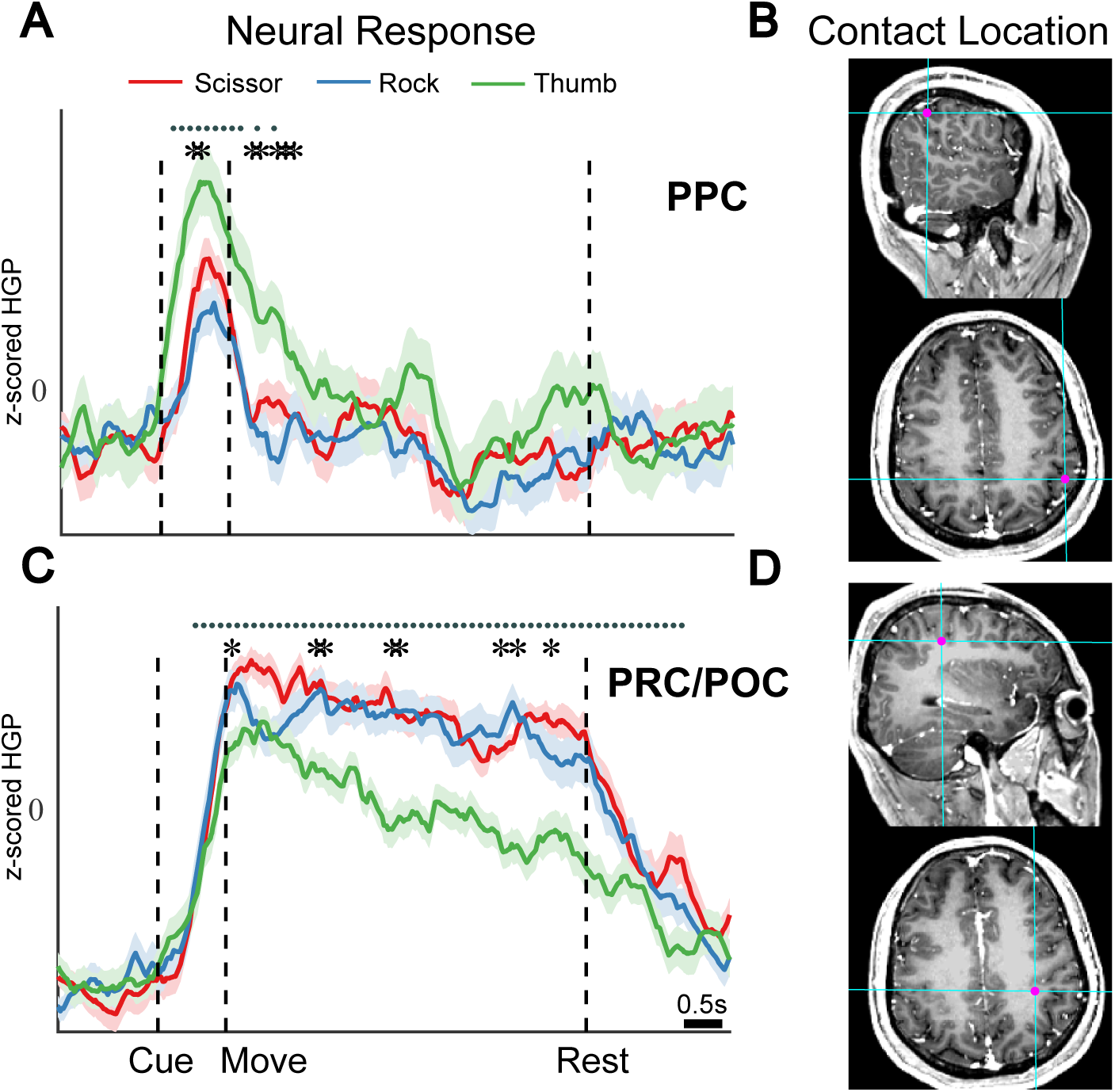
The neural response of two exemplary electrodes located in different ROIs from subject S1. **(A)** The z-scored HGP of an electrode located in PPC of left hemisphere during the task. The solid line indicates the mean HGP across 20 trials of each gesture. The shading area indicates the standard error. The dot at the top denotes the significantly activated time windows when 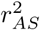 is significant (*p <* 0.001, FDR corrected) and the asterisk above the lines denotes the significantly selective windows when 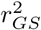 is significant (*p <* 0.05, FDR corrected). **(B)** The contact location of the PPC electrode in (A). The electrode is rendered as the magenta dot on sagittal and axial MRI images respectively. The Talairach coordinate of this contact is [-49 −37 47]. **(C)** The z-scored HGP of an electrode located near central sulcus of left hemisphere during the task. Lines and symbols in the figure use the same conventions as (A). **(D)** The contact location of the primary sensorimotor electrode shown in (C). Same convention as (B). The Talairach coordinate of this electrode is [-31 −25 41].

Noticing that the activation in different ROIs were not synchronized in time (figure 4), we further investigated whether there existed a temporal activation sequence in such a visuomotor task. To finalize this, we pooled the first activation time (FAT, Sec. 2.6.3) of all significantly activated electrodes within each ROI in table 2. As a result, FAT value from 147 electrodes in PRC, 68 electrodes in POC and 265 electrodes in PPC were obtained. Within these ROIs, the median FAT was 250 ms for PPC, 400 ms for PRC and 650 ms for POC after *Cue* respectively. Besides, when comparing with average *Move* time (564 ms, Sec. 2.5), the FAT was [-0.35, −0.25] s, [-0.15, −0.15] s and [-0.15, 0.05] s (95% confidence interval, Sec. 2.6.3) for PPC, PRC and POC respectively (figure 5). The activation of ROIs were significantly sequential along time course, where PPC activated first, PRC second and POC last (*p <* 0.001 between PPC and PRC, *p* = 0.06 between PRC and POC, one side Wilcoxon Ranksum tested after Bonferroni correction). The first selectivity time was not compared due to the relative small sample set (19 electrodes in PRC, 15 electrodes in POC and 8 electrodes in PPC).

**Figure 5:**
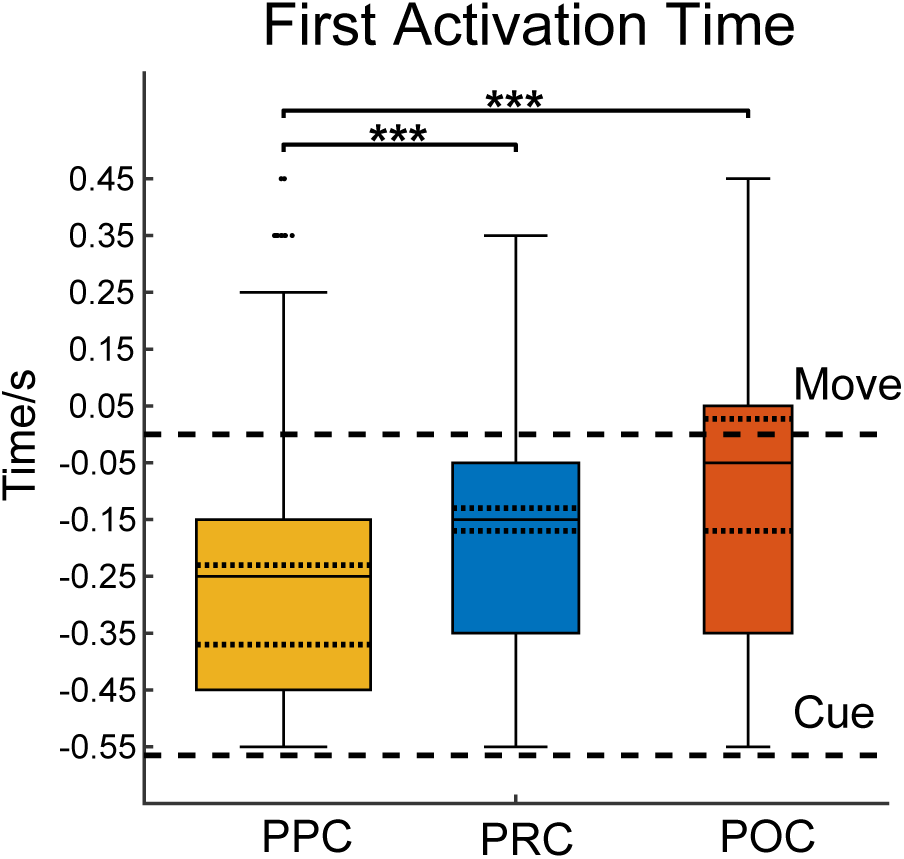
First activated time (FAT) of each ROI. The dash line inside each box indicates the 95% confidence interval by bootstrap (Sec. 2.6.3). Inside each box, the black solid line denotes the median FAT (*q*_2_). The lower and upper boundary of the box denotes the 25% (*q*_1_) and 75% (*q*_3_) percentile. The width of each box is scaled to the number of samples (i.e., activated electrodes). The whisker of each box extends most to *q*_3_ + 1.5(*q*_3_−*q*_1_). The black dot outside whisker denotes the outliers. The long dash lines outside box are the median *Move* time (upper) and *Cue* time (lower). *** indicates *p <* 0.001 (one side Wilcoxon Ranksum tested after Bonferroni correction). *p* = 0.06 for the test between PRC and POC.

### 3.3. Gesture Decoding from ROIs

To determine if PPC could assist PRC and POC for gesture decoding, we first identified the three subjects who had selective electrodes in both PRC/POC and PPC within [-0.5, 0.5] s around *Move*. Then only the selective electrodes from these identified subjects were used for further analysis in order to make fair comparison. For the identified subject, the number of electrodes used for decoding was: 1/1/5 (S1), 2/5/1 (S3), and 2/1/2 (S4) in order of PRC/POC/PPC. The obtained DAs was shown in figure 6. As could be seen from figure 6A, combining PPC throughout *Move* as spatial feature improved the DA by 5% on average compared to only using primary sensorimotor cortex, which indicated that PPC could provide effective and complementary information to primary sensorimotor cortex for gesture decoding. To account for the factor that such improvement was due to higher feature dimension on condition of adding PPC, we randomized the PPC feature across trials and repeated the decoding process together with normal features from primary sensorimotor cortex to yield the “With Random PPC” accuracy. The results demonstrated that the addition of random PPC was unable to improve the DA (figure 6A), confirming that the PPC contained useful visuomotor information that can assist the gesture decoding.

**Figure 6:**
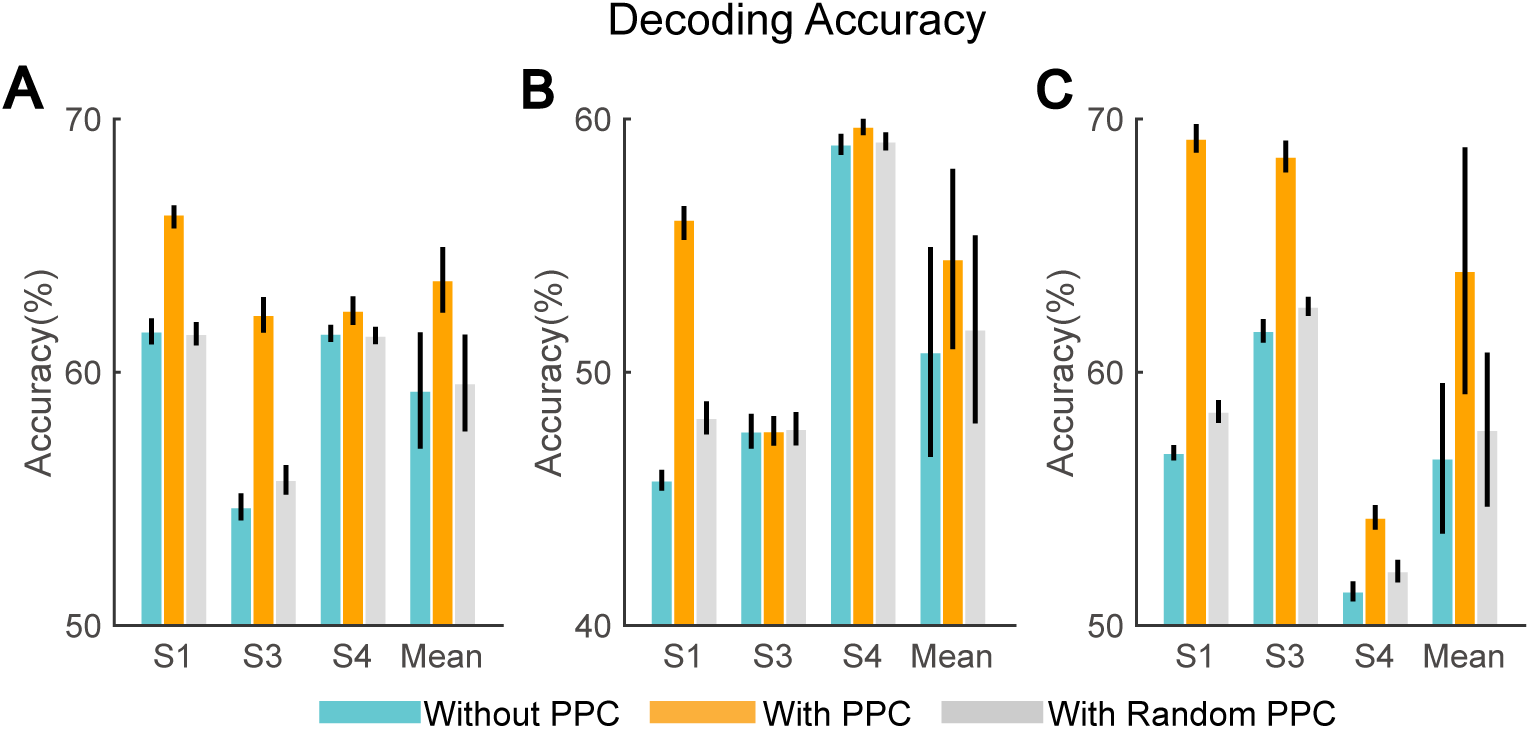
Decoding performance with and without PPC. **(A/B/C)** DA obtained using temporal features drawn from different time period: (A) [-0.5, 0.5] s, (B) [-0.5, 0] s, and (C) [0 0.5] s. *Blue*: DA computed by using electrodes located at PRC and POC only; *Orange*: similar to *Blue* but PPC is also included; *Gray*: similar to *Orange* but PPC was randomized across trials and included. For each subject, the errorbars represent the standard error over fOFS (see Sec. 2.6.4). For mean accuracy over the three subjects, the errorbars represent the standard error over the final DA of the three subjects. For A, B, and C, Wilcoxon signed-rank test between with and without PPC shows no significance. *p* = 0.125 for all. No significance between without and with randomized PPC as well.

Moreover, we further asked whether PPC could assist prediction decoding both in advance of and later than *Move*. To verify it, we implemented the same decoding process using selective electrodes from PPC and primary sensorimotor cortex, but changing the temporal range of the feature vector from [-0.5, 0.5] s to [-0.5, 0] s and [0 0.5] s around *Move*, respectively (Sec. 2.6.4). The results were shown in figure 6B and C. With PPC improved the DA by 3.6% before *Move* and 8% after *Move* on average while with random PPC only slightly improved by about 1%. Even though increases of DA could been observed from all the subjects (largest for S1, slight for S3 and S4) when introducing the features from PPC, no significance was achieved (*p >* 0.05, figure 6).

## 4. Discussion

In current study, we evaluated whether human PPC encoded neural information related with hand gestures, and moreover, investigated for the first time whether PPC together with primary sensorimotor cortex (PRC and POC) could enhance the hand gesture decoding using SEEG recordings from 25 human subjects participating a hand gesture experiment. In our evaluations, we proposed four performance indices and aimed to answer a series of relevant critical questions: 1) in what extent PPC is involved to the task; 2) if there exists temporal activation sequence between PPC and primary sensorimotor cortex during the visuomotor task; 3) whether iEEG recordings from PPC contains the fine gestures information and can subsequently improve decoding performance.

### 4.1. Temporal activation sequence

Incorporating LFP recordings across multiple subjects, we have found that a majority of electrodes located in PPC, PRC and POC are activated with a sequence in terms of HGP, where PPC activates first, PRC second and POC last (figure 5). Two other human iEEG studies (Coon et al., 2016; Coon and Schalk, 2016), which are not directly addressing the same question with this work but also using HGP and conducting similar behavior task, present similar results. In Coon et al. (2016, Fig.2), three ECoG electrodes located at PPC, PRC and POC are sequentially activated around 250 ms, 400 ms and 590 ms after *Cue* with EMG onset at 576 ms, similar to 310 ms, 400 ms, 510 ms after *Cue* with EMG onset at 564 ms in current study (figure 5).

Considering that - 1) PPC is viewed as an earlier motor intention formation cortex (Andersen and Buneo, 2002); 2) PRC and POC are responsible for the motor execution and somatic sensations processing respectively, current result is in agreement with a classic cognitive view about visuomotor task that perception, cognition, motor control and somatosensory feedback sequentially occurs. Notably, there are also studies report different results conflicting with the classic view. For example, some neurons in PPC keep active during the whole Cue-Delay-Move process (Klaes et al., 2015; Aflalo et al., 2015), which is not compatible with any sole role in a sequential perspective. Ledberg et al. (2007) demonstrate that an early evoked potential occurs in primary motor, premotor and prefrontal cortex immediately after visual cortex in a visuomotor task, thereby showing that primary motor cortex activates earlier than PPC without early stimulus-related processing. An emerging perspective to reconcile these evidences holds that perception, cognition, motor and feedback may not sequentially occurs and does not map cortex and function in a one-to-one manner (Cisek and Kalaska, 2010). In this view, the visuomotor task involves a multi-stage forward and backward activation among ROIs. However, this view does not exclude the large-scale temporal sequence presented by current results, which depends on how many neuron populations are involved in each stage. Therefore, it is possible that in a finer scale, a multi-stage forward and backward activation sequence exists, while in a large-scale, each ROI has a relative temporal sequence.

Notably, even though POC activates lastly in sequence, the detected activation of POC in current work may represent more than somatosensory feedback after motor execution, since the average activation time of POC electrodes in this work is slightly ahead of EMG onset (figure 5), indicating that some neurons in POC initiate firing before movement onset. The early activation of somatosensory cortex may suggest that some neurons in this area encode sensory information in anticipation of movements (Sun et al., 2015).

### 4.2. PPC assists gesture decoding

The LFP results in this work show that human PPC is selective to hand gestures (figure 3), verifying our hypothesis that human PPC iEEG recording contains gesture related information, which is consistent with previous monkey LFP studies and human spike studies (Asher et al., 2007; Baumann et al., 2009; Klaes et al., 2015). In this work, we first observed that the electrodes were widely activated among each ROI (table 2, figure 3). This finding is also in agreement with reported ECoG studies (Caruana et al., 2017; Kubanek et al., 2009) and thus supports a widespread visuomotor circuit shown by the fMRI study (Króliczak et al., 2016). Among the largely activated electrodes, only a few electrodes in PPC present selectivity without high *r*_*GS*_ while most activated electrodes are not selective, which is much less than 37% shown in Asher et al. (2007). Several reasons may account for this. First, the hand shape selective area may indeed occupy a small area of PPC, most likely AIP. Although several fMRI studies indicate the hand shape selective area includes beyond AIP such as SMG, the whole IPS, and SPL (Fabbri et al., 2016; Króliczak et al., 2016; Ariani et al., 2015), AIP is consistently found to take part in the special encoding of hand shape (Sakata et al., 1995; Binkofski et al., 1998; Klaes et al., 2015), which has been demonstrated in this work as well (figure 3). Therefore, since SEEG electrodes are randomly distributed within the whole PPC, it is reasonable that only a small percentage of them presented gesture selective. But because of the comparatively limited number of implanted electrodes relative to the large PPC, we hesitate to conclude the exact location which still needs more data to answer. Secondly, it may come from the blurring effect of LFP on a inherently less prominent encoding of hand shape in PPC. Either fMRI studies or macaque spiking studies indicate that PPC is relatively less informative for classification of grip types compared to primary sensorimotor cortex or premotor cortex (Fabbri et al., 2016; Townsend et al., 2011). A weaker neuronal encoding is easier to be blurred in the wide listening area of LFP (Buzsáki et al., 2012). This conjecture is in coincide with the result in Asher et al. (2007) that the selective LFP contains less information about grip type than spiking activity when they are recorded in the same area of PPC, suggesting a blurred effect. Therefore, it may underline that many electrodes in PPC are activated but not selective (figure 3). The small group of selective electrodes in PPC suggests that the hand shape selective sites may be spatially sparse within PPC.

Nevertheless, the small group of PPC selective electrodes still provides effective information for classification and such information is complementary to the primary sensorimotor area (figure 6). For instance, the PPC electrodes in figure 4A and supplementary figure 2A respond more actively to more precise gestures like thumb and scissor than whole hand gesture like rock, and hence may indicate the different gesture encoding and this finding is in accordance with previous fMRI study (Begliomini et al., 2007). Moreover, these PPC electrodes also respond differently from the primary sensorimotor electrodes, thus collectively improving the decoding accuracy (figure 6A) by 5%. The non-significance may be due to the small sample size (*p* = 0.125, *n* = 3). As for pre-movement and post-movement decoding (figure 6B and C), we find that both of them present enhancement of performance in an average view while post-movement holds higher enhancement, where these enhancement are also in line with the phenomenon that the selectivity is found both before and after *Move* (figure 4 and supplementary figure 2). In addition, the HGP response after *Move* shows smaller variance than before, which leads to less feature overlap among gestures and thus contribute to the higher post-movement classification performance. Moreover, by identifying the pre-and post-movement decoding performance enhancement from PPC, our results extend the finding in Asher et al. (2007), which revealed that selectivity of movement-evoked LFP in macaque PPC appeared only after movement onset. Furthermore, the implication of this difference between pre- and post-movement will be discussed in next section (Sec. 4.3).

### 4.3. PPC: Visual or Motor?

The role of PPC especially AIP in the visuomotor task is more related to visual shape of objects or the hand shape is still in debate. The common view from previous reports is that it contains visual-dominant, motor-dominant and visuomotor neurons (Mountcastle et al., 1975; Sakata et al., 1995; Murata et al., 2000; Baumann et al., 2009; Townsend et al., 2011; Klaes et al., 2015). Specifically, in a typical Cue-Delay-Go grasp task, the visual-dominant neurons only respond during Cue phase while motor-dominant neurons only respond during Delay or Go phase even in darkness, where the visuomotor neurons respond during both phases (Murata et al., 2000). In a non-object related gesture task including scissor, rock, and paper picture, there still exist similar visual-dominant and motor-dominant neurons (Klaes et al., 2015). Therefore, in the view of the different response phases, it is believed that PPC encodes visual and motor information sequentially. But due to the intrinsic overlap between object shape and hand shape in the studies above, it is still unclear that the response during each phase is encoding movement details or object property. To address this question, a recent fMRI study on human, which orthogonally changed the shape of objects and the hand shape required to grasp, revealed that PPC during grasping encodes not only the grip type but also the number of digits used, also object elonganation and shape, where the previous two aspects are corresponding to hand shape details while the latter two aspects to object property, thus supporting the mixing encoding view. Taken together, PPC may encode visual properties of object, transform these sensory information to motor intention, and hold the visual object properties while receiving the feedback of motor signals from ventral premotor area during hand movement execution (Jeannerod et al., 1995; Schaffelhofer and Scherberger, 2016).

Therefore, the response of PPC in current study may reflect this mixture encoding scheme. Specifically, the HGP results in figure 4 and decoding results in figure 6 suggest that the encoding of hand gesture exist both before and after movement onset. Considering that LFP is the summation of many different neural activities, its encoding components before movement likely come from the visual-dominant neurons, while that after movement come from motor-dominant neurons. This interpretation also explains the higher enhancement of decoding accuracy after *Move*, because the exact encoding of hand movement is more likely to be stronger represented in motor neurons than in visual neurons. On this basis, the post-movement HGP in PPC may reflect the hand shape rather than visual stimuli, implicating its potential for BMI application. To thoroughly evaluate this speculation, a new task in dark environment including delay phase and orthogonality between sensory cue and hand gesture will be designed and conducted under a larger subject population in our further study.

## 5. Conclusions

In this work, we assessed the human PPC response during a visually-cued hand gesture task and investigated its potential benefits to conventional primary sensorimotor cortex-based gesture decoding using SEEG recordings for the first time. Using dataset from 25 subjects, we observed significant activation for all three areas during the task and they tended to activate in a temporal sequence: PPC first, PRC next, and POC last. The study demonstrates a spatially wide and temporally early involvement of PPC in a visuomotor task, supporting the notion that PPC plays an important role in early visuomotor processing. Moreover, we identified that some of the activated electrodes within primary sensorimotor cortex and a minority within left PPC presented selectivity across different gestures, implying that fine hand neural encoding information in PPC is likely spatially sparse. Finally, we found that gesture decoding performance could be improved by using SEEG signals from both PPC and primary sensorimotor cortex and the results were valid within the temporal range either before and/or after movement. In summary, our results suggest that human PPC encodes specific information about fine hand movement which is complementary to that of primary sensorimotor cortex, potentially providing a new signal source that will benefit further iEEG-based BMI applications.

## Acknowledgments

This work was supported by grants from the National Natural Science Foundation of China (No. 91848112, No. 61761166006), the Shanghai Municipal Commission of Health and Family Planning (No. 2017ZZ01006), and the Shanghai Municipal Science and Technology Major Project (No. 2018SHZDZX03).

## Appendix A.

### Abbreviations

Alphabetical list of abbreviations used in this paper.

AIP: Anterior Intraparietal Area
AS: Activation Strength
BMI: Brain-Machine Interface
DA: Decoding Accuracy
ECoG: Electrocorticography
EMG: Electromyography
FAT: First Activation Time
fMRI: Functional Magnetic Resonance Imaging
GS: Gesture Selectivity
iEEG: Intracranial Electroencephalography
IPS: Intraparietal Sulcus
LFP: Local Field Potential
MNI: Montreal Neurological Institute
POC: Postcentral Cortex
PPC: Posterior Parietal Cortex
PRC: Precentral Cortex
ROI: Region of Interest
SEEG: Stereo-Electroencephalography
SMG: Supramarginal Gyrus
SPL: Superior Parietal Lobe

## Appendix B. Supplementary Material

**Supplementary Fig. 1:**
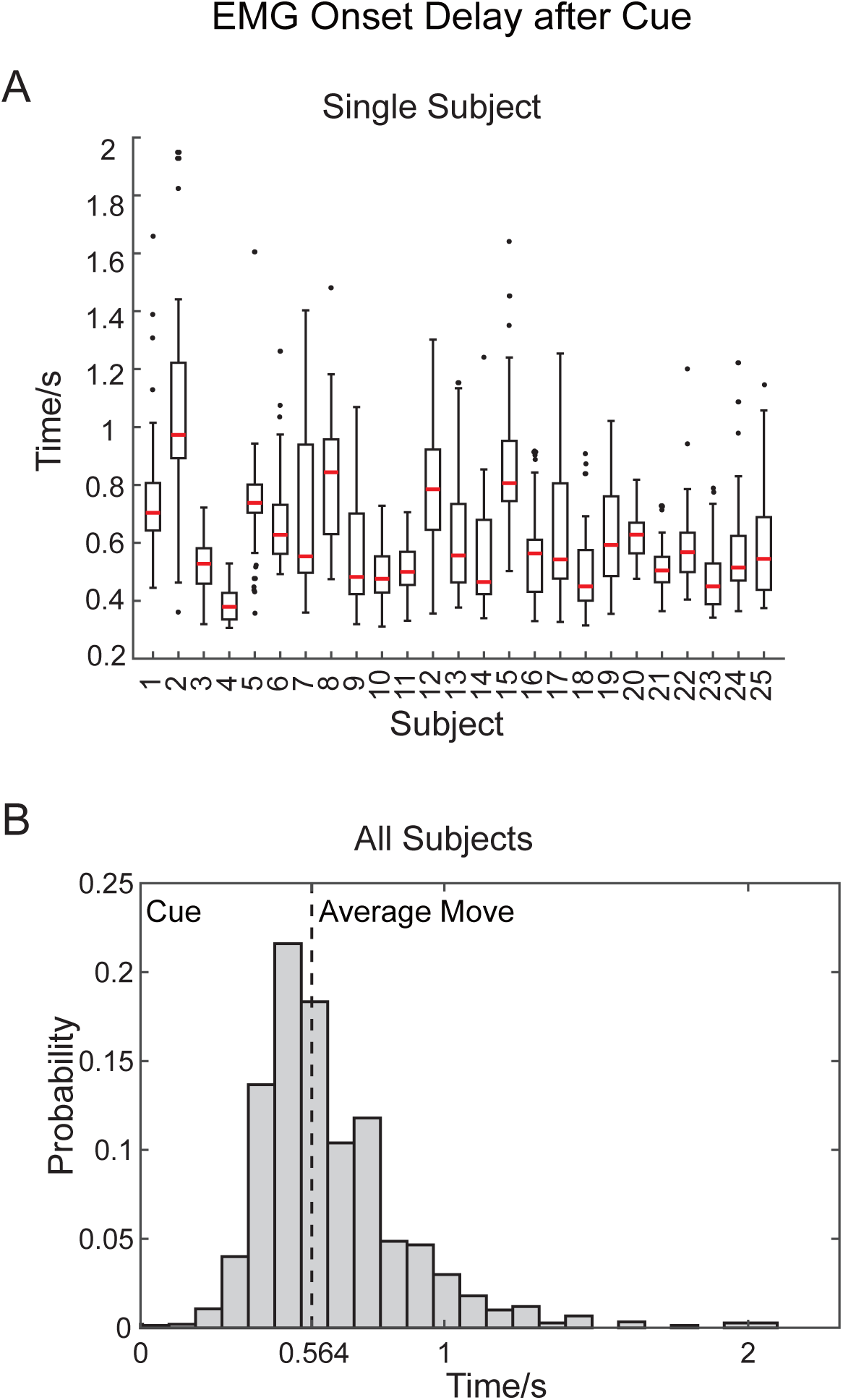
**A** Boxplot of EMG onset delay after *Cue* of each subject. The red line denotes the median time. The upper and lower boundary of box denotes 25% and 75% percentile. The black dots denotes the outlier. **B** The average delay over all subjects is 564 ms. 564 ms after cue appears, the subject started to move their hand.

**Supplementary Fig. 2:**
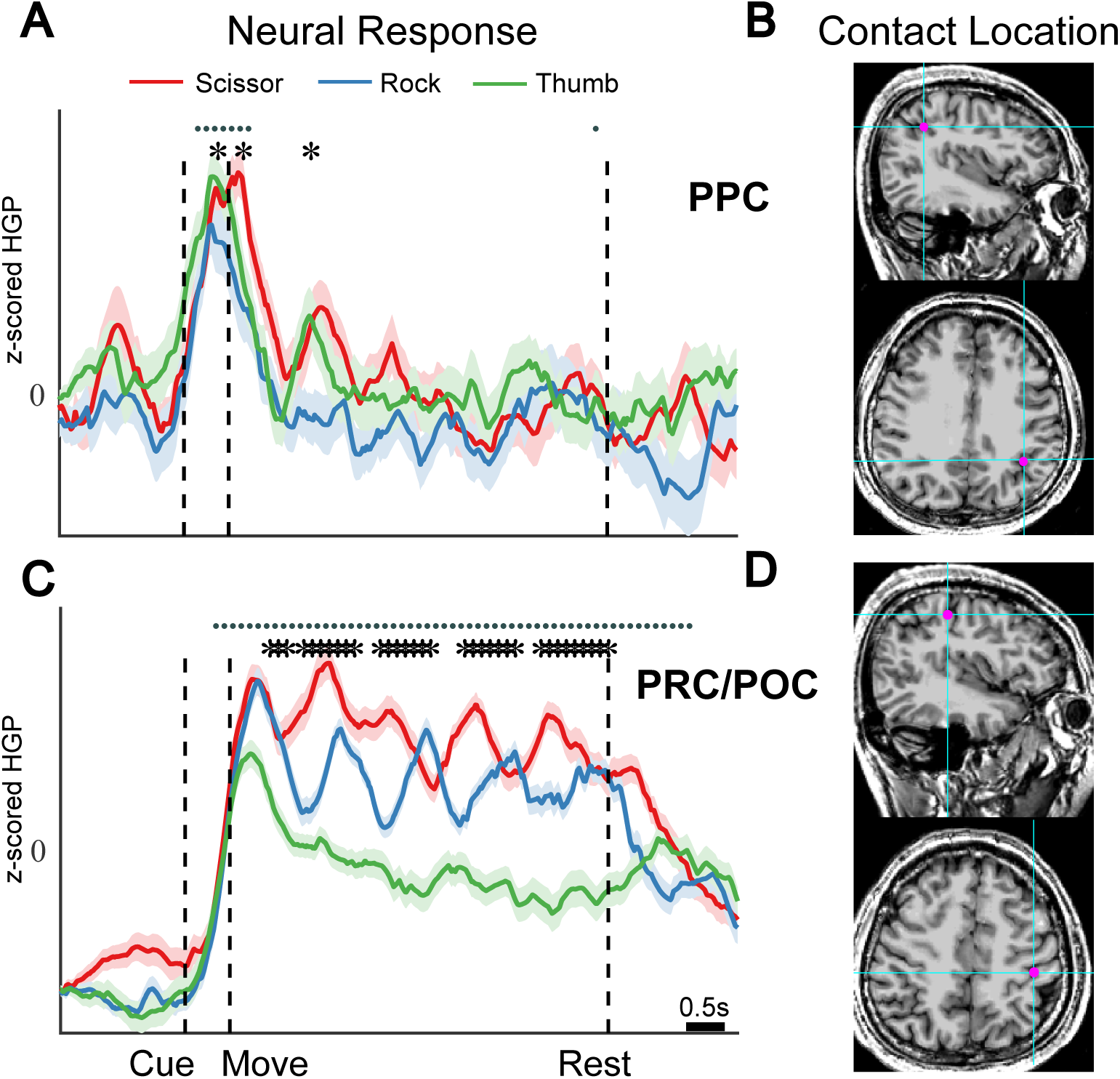
Similar convention as figure 4 but electrodes of S3. The Talairich coordinate in (C) is[-35 −41 37], and in (D) is [-36 −19 46].

